# MechPPI: Binding Mechanism-based Machine-Learning tool for Predicting Protein-Protein Binding Affinity Changes Upon Mutations

**DOI:** 10.1101/2023.10.26.564257

**Authors:** Yangying Liu, Grant Armstrong, Justin Tam, Brian Y. Chen

## Abstract

Protein-protein interactions are essential for various biological processes, including signal transduction, metabolism, vesicle transport, and mitogenic processes. It’s crucial to consider them within the context of their interactions with other proteins to understand protein function. Mutations in proteins can affect their binding affinity to partner proteins by introducing various effects, such as changes in hydrophobic regions, electrostatic interactions, or hydrogen bonds. Assessing the impact of mutations on protein interactions can have implications for disease susceptibility and drug efficacy. Understanding the impact of mutations on protein-protein interactions and predicting binding affinity changes computationally can benefit both basic biology and drug development. Different computational methods offer varying levels of accuracy and efficiency, and the choice of method depends on the specific research goals and available resources. We developed MechPPI, a tool that can use potential mechanism features underlying mutation to predict the binding affinity change upon mutation. We showed MechPPI can accurately predict binding affinity change upon a single mutation, and results demonstrate the potential of MechPPI as a powerful and useful computational tool in protein design and engineering.

## 1 Introduction

Protein-protein interactions (PPIs) play a vital role in various fundamental biological processes, including signal transduction, metabolism, vesicle transport, and mitogenic processes [1]. Therefore, in order to understand the function of the proteins, they must be considered within the context of other interacting proteins. Some protein interactions within a cell are theoretically possible, however, only a subset actually results in functional complexes and assemblies. The binding energy of these protein interactions plays a crucial role in determining the stability and conditions for forming these complexes [2]. Hence, a complete comprehension of cellular processes requires not only knowledge of all possible Protein-protein interactions but also quantitative insight into the structure and stability of the formed complexes [2–4].

The strength of the interaction between a protein and its binding partner is the binding affinity and this thermodynamics value influences the stability of protein complexes. The equilibrium dissociation constant (*K*_*d*_) or Gibbs free energy (ΔG, also called the binding free energy), which can be derived from the *K*_*d*_, can be commonly used to quantify binding affinity of protein-protein interactions. The binding affinity of complexes can be affected by mutations in proteins, because mutations introduce various types of effects, which include reduction in a hydrophobic region, decrease in electrostatic interactions, loss of hydrogen bond, and so on, which can modulate or even disrupt the binding to partner proteins, further leading to the dysfunctionality of proteins in a cell [5–7]. Hence, assessing the impact of mutations on how proteins interact with each other can shed light on disease susceptibility and drug efficacy [7, 8]. The impact of a mutation on the stability of protein-protein interactions can be measured by the difference in binding free energy of the mutated and wild-type complexes. This difference in ΔG between mutant and wildtype protein complexes is defined as the binding affinity change (ΔΔG) caused by mutations:

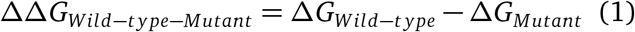

Traditional methods for studying the binding affinity of protein complexes are through experimental techniques, which are expensive and time-consuming. To solve the above-mentioned limitations of experimental measurement, computational methods for predicting binding affinity upon mutations are developed, including molecular dynamics simulations, which simulate the difference of free energy between the wild type and the mutant of the system, this type of method provides reliable and acurate results but have high requirement for computational resources, limit their prediction of large-scale variants; empirical energy functions, using empirical force field or statistical potentials to approximate the energy of interactions between atoms in a molecule to calculate the free-energy changes, FoldX [9] and Rosetta [10] are two representative methods that combine empirical energies as linear functions to estimate ΔΔG, this type of method speeds up the prediction but suffers from insufficient conformational sampling and generalization performance remained poor; machine learning methods, fit the experimental data using intricate design features of the changes in structures [9, 11, 12].

With the accumulation of data and growing volume of experimental data in SKEMPI 2.0 dataset, machine-learning techniques can be developed to predict the correlation between a mutation and the subsequent change in binding affinity in a more acurate and efficient way. For example, mCSM-PPI2 [13] uses the Extra Tree method to do prediction with features including atomic distance patterns to indicate environmental information surrounding mutation sites, and other six physicochemical features for both wild- and mutant-type residues as input for a regression model. Muta-Bind2 [14] used seven features, including interactions of proteins with the solvent, evolutionary conservation, and thermodynamic stability of complexes for the prediction model. GeoPPI used an autoencoder to extract deep geometric representations of protein complexes from protein structure and integrated with gradient-boosting trees for prediction, which achieved state-of-the-art performance on the SKEMPI 2.0 dataset. Some of the existing machine-learning methods use physical quantities as features, which is time-consuming. In addition, some input features are mostly manually engineered from protein structures and limit their predictive generalization across various protein structures. Most importantly, most methods used features only include structural information and some physicochemical information about key residues, which might not be enough for predicting binding affinity change upon mutation. Mutations not only introduce structure and physicochemical change for protein complexes and key residues but also introduce changes in the binding mechanisms, such as reduction in a hydrophobic region, decrease in electrostatic interactions, loss of hydrogen bond, and so on. These binding mechanisms directly contribute to the binding affinity of protein complexes and provide more valid information for prediction.

In this work, we aim to develop a method that not only provides fast and acurate predictions but also includes acurate change of binding mechanism information upon mutation, such as electrostatic complementarity, hydrogen bonding, and steric hindrance. We propose a novel method, called MechPPI, to predict the effects of mutations on the binding affinity. Compared with traditional methods, we feature the following advantages of MechPPI: 1) MechPPI provides a more accurate way to predict binding affinity with less error. 2) MechPPI has provided insight into binding mechanisms that are the driving factors for the binding process across different protein complexes upon mutation.

## 2 MATERIALS AND METHODS

### 2.1 Datasets

Datasets used to construct the model were from the SKEMPI v.2.0 database [15]. SKEMPI v.2.0 is a comprehensive database that includes information about mutations, kinetic and thermodynamics in protein protein interactions (PPIs) that have been resolved structurally. It is the most complete source for experimentally measured binding affinities of wild-type and mutated protein complexes, including 7,085 documented mutations, it’s currently served as a commonly used resource for exploring the structural and energetic consequences of mutations on PPIs. This manual curation ensures the accuracy and reliability of the information, making it a vital tool for studies in protein engineering, molecular dynamics, and drug design.

The most important thermodynamic parameter in the database is dissociation constants (*K*_*d*_), which are used to represent affinity. Here, we calculated the change in free energy (ΔG) through the formula: ΔG = RTln(*K*_*d*_), where R is the ideal gas constant(8.314 J *mol*^*-*1^*K*^*-*1^) and T is the absolute temperature. Both mutants’ and wild-type free energy were calculated and get the difference between them as binding affinity change(ΔΔ*G*, ΔΔ*G*_*wild*–*type*–*Mutant*_ = Δ*G*_*Wild*–*type*_ − Δ*G*_*Mutant*_). and Four widely used subsets of SKEMPI v.2.0 were used as benchmark datasets, namely S1131, S4169, and S8338. S1131 was a subset of 1,131 nonredundant interface single-point mutations. S4169 includes selected 4,169 single-point mutations derived from SKEMPI v.2.0. S8338 includes 8,338 data points, was derived from S4169 plus all hypothesized reverse mutations, setting the reverse mutation free energy changes to the negative values of the original free energy changes.

### 2.2 MechPPI Overview

MechPPI is a machine learning-based method that uses both binding mechanisms and structural features of protein complexes to predict the binding affinity upon mutations. Our methods are different from other methods, there are four main feature groups included in our model, and they are related to potential binding mechanisms under mutation of the protein complexes. A machine learning method gradient-boosting tree (GBT, excelling in avoiding overfitting) is used to leverage these features for exploring the effects of mutations on the binding affinity of protein complexes.

### 2.3 Binding mechanism based features

#### 2.3.1 Electrostatic features

Electrostatic interactions play a key role in stabilizing protein complexes, these interactions are attractive forces between oppositely charged regions and repulsive forces between like-charged regions, which contribute significantly to binding affinity. When molecules or molecular surfaces exhibit electrostatic complementarity, it means that regions with positive charges on one molecule align with regions of negative charges on the other molecule, promoting strong attractive interactions. Hence the degree of electrostatic complementarity between binding partners correlates well with the binding affinities. Here, we quantitatively measure the electrostatic complementarity degree of the protein complexes through solid volume representations. We identify regions by representing electrostatic isopotentials with volumetric solids generated by genSurf. For example, we can get the three-dimensional contour for electrostatic isopotentials at +1 (kT/e) and use genSurf to get a volumetric representation for the electrostatic isopotentials region of one protein. Similarly, we can also get a volumetric representation of electrostatic isopotentials at -1 (kT/e) for its binding partner. Then, the intersection volume of the solid representation of electrostatic isopotentials between protein at +1 (kT/e) and its binding partner at -1 (kT/e) for protein complexes reflects electrostatic complementarity to some extent.

Therefore, we can obtain the degree of electrostatic complementarity change between wild-type and mutant protein complexes by acquiring the difference in that volume between wild-type and mutant protein complexes. Here, we get four types of solid volumetric intersection for electrostatic isopotentials of protein complexes: 1) the intersection between the volume of electrostatic isopotentials at - 1 (kT/e) for one protein and the volume of electrostatic isopotentials at +1 (kT/e) for its binding partner, 2) the intersection between the volume of electrostatic isopotentials at +1 (kT/e) for one protein and the volume of electrostatic isopotentials at v-1 (kT/e) for its binding partner, 3) the intersection between the volume of electrostatic isopotentials at +5 (kT/e) for one protein and the volume of electrostatic isopotentials at -5 (kT/e) for its binding partner, 4) the intersection between the volume of electrostatic isopotentials at -5 (kT/e) for one protein and the volume of electrostatic isopotentials at +5 (kT/e) for its binding partner. We calculated all four types of electrostatic isopotentials volume intersection for the wild-type protein complexes and mutant protein complexes, as well as the difference between wild-type protein complexes intersection and mutant protein complexes intersection. All intersection volumes are regarded as electrostatic complementarity feature group and provide electrostatic interaction mechanisms for predicting the effect of mutation on protein complexes.

#### 2.3.2 Hydrogen Bonding features

In this study, we investigated the intricate network of hydrogen bonds at the interface between two interacting proteins (protein-protein interaction, PPI) and their susceptibility to mutations. Hydrogen bonds play a pivotal role in stabilizing protein complexes and are essential for the specificity and strength of protein interactions. Mutations in protein complexes can disrupt these hydrogen bond networks, leading to alterations in binding mechanisms and affinity. Importantly, even mutations occurring at sites distant from the PPI interface can induce changes in the hydrogen bond network, illustrating the long-range effects of amino acid substitutions on protein-protein interactions.

To quantify the hydrogen bond interactions, we tracked four distinct types of hydrogen bonds: 1) backbone-backbone, 2) backbone-sidechain, 3) sidechain-sidechain, and 4) sidechain-backbone interactions. This comprehensive analysis allowed us to create a measurable “ontology” for each PPI complex, characterizing the intricate hydrogen bond landscape within these complexes.

To assess the impact of mutations on the hydrogen bond network, we compared the network in the mutant (MT) complexes to that in the wild-type (WT) complexes at two scales. The first scale encompassed the entire interfacial hydrogen bond network, which we refer to as the “supergraph.” The supergraph included all hydrogen bonds between protein 1 and its binding partner protein, providing a holistic view of the intermolecular interactions. The second scale focused exclusively on the subnetwork of hydrogen bonds involving the mutated amino acid. This subnetwork, referred to as the “sub-graph,” was a subset of the supergraph and offered insight into the local hydrogen bond ontology near the mutated residue. Additionally, we quantified the “offsite effects” as the difference between the super-graph ontology change and the subgraph ontology change. Offsite effects highlighted alterations in hydrogen bonding interactions that were non-local to the mutated amino acid but still within the PPI interface. These hydrogen bonding features serve as valuable indicators for capturing the changes in binding mechanisms induced by mutations. They play a pivotal role in shaping the binding process across various protein complexes. Importantly, the use of these hydrogen bonding features is more efficient than existing methods that rely on computationally intensive biophysical simulations. This efficiency stems from the fact that these features can be measured directly from the single-frame X-ray crystallographic poses, providing a practical and informative tool for studying the effects of mutations on protein-protein interactions. We further employ volumetric spherical cones as a fundamental geometric representation of hydrogen bond interactions. These spherical cones are generated based on well-documented angle and distance tolerances associated with hydrogen bonds. Each hydrogen bond interaction involves both an acceptor and a donor atom, and for each of these atoms, a spherical cone is constructed. The crucial criterion for identifying an intersection surface between these cones is that they must be oriented to face each other and possess overlapping radii. When these conditions are met, an intersection surface is formed. The union of all intersecting spherical cones results in the creation of a volumetric intersection region, the volume of which can be precisely measured as a continuous value. This volumetric intersection region serves as a key feature in our predictive model, allowing us to quantitatively assess the extent of hydrogen bond interactions within protein complexes and providing valuable insights into the structural basis of protein-protein interactions.

#### 2.3.3 Steric features

Steric hindrance can have significant effects on the geometry of protein and protein complexes. It can influence bond angles, bond lengths, and the overall three-dimensional shape of a molecule. When atoms or groups of atoms are too close to each other, they experience steric repulsion, which can lead to distorted bond angles and strained conformations, further influencing binding affinity. Mutation might introduce steric effects on the protein complexes, this influence might be diverse and dependent on different protein complexes and mutation types. For example, mutation of a small residue to a larger one could cause steric clashes, while the reverse could create holes and minimize steric effects, both cases could affect local structural conformation and further affect the binding affinity of protein complexes. Hence, include the amino acid volume of wild-type protein complexes and the amino acid volume of mutant protein complexes. However, only knowing amino acid volume can not provide enough steric information for protein complexes. Point mutations in different locations are of different importance. When amino acid substitution is from a small one to a larger one, the protein noncore accommodates changes in amino acid side chains more easily compared to the protein core, and thus changes on the surface are less likely to be lethal for protein integrity and stability [5]. Therefore, in order to include point mutation location information, we calculate the nearest pairwise atomic distance of the target amino acid to the opposite chain. If the distance is less than 5 A, we can regard it as an interfacial amino acid, otherwise, it’s a non-interfacial amino acid. Interfacial amino acids can be regarded as buried amino acids that are in general predicted better than exposed ones. Even though amino acid volume and the nearest pairwise atomic distance provide steric information, these are relatively coarse features. Besides, we use molecular surface intersection volume generated by genSurf to provide a finer measurement for the steric effect of the interface of protein complexes. Firstly, we get the volume of the molecular surface for one protein and its binding partner, then use genSurf to get the difference between them as molecular surface intersection volume. For all these three types of steric features, we included amino acid volume, nearest atomic distance, molecular-surface intersection for wild-type protein complexes, mutant protein complexes, as well as differences between wild-type protein complexes and mutant protein complexes.

#### 2.3.4 Hydrophobic features

Hydrophobic interactions occur because some amino acid side chains are hydrophobic, meaning they repel water molecules. These hydrophobic side chains tend to cluster together in the interior of a protein, away from the surrounding aqueous (water-based) environment. Here, according to Kyte-Doolitle scale [16], we evaluate the hydropathy of target amino acid and mutated amino acid, as well as the difference between them. We also put the nearest atomic distance into hydrophobic feature group, because hydrophobic areas tend to be located in the interior of the protein, away from the surrounding aqueous environment. While hydrophobic residues are predominantly found in the core, proteins may also have specific hydrophobic regions on their surfaces. These regions can be involved in protein-protein interactions, where one protein binds to another. In such cases, hydrophobic interactions between these regions can play a role in forming and stabilizing protein complexes.

## 3 RESULTS

### 3.1 MechPPI acurately yields PPI ΔΔG

To comprehensively compare MechPPI with other existing advanced approaches, we considered three single mutation benchmark datasets, including single-point mutations in the, non-redundant interface single-point mutations in the SKEMPI dataset (S1131) [17], single-point mutations in the SKEMPI2 dataset (S4169) [13], and single-point mutations in the SKEMPI2 balanced dataset (s8338) [13].

#### 3.1.1 10 fold cross validation

Based on the above three datasets, firstly we performed the 10-fold cross-validation splitting data based on mutation sample points and used mean Pearson’s correlation coefficient (*R*_*p*_) and Rooted mean square error(RMSE) as two main evaluation metrics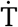herefore, all mutation sample points in each dataset will be split into 10 folds for the 10-fold cross-validation. Then we compared them with seven baseline methods for PPI ΔΔ*G* predictions, including TopNetTree, GeoPPI, mCSM-PPI2, DGCddG, MutaBind2, FoldX, and BeAtMuSic. For set S4169, we obtained a high *R*_*p*_ of 0.783 and lowest RMSE of 1.11 *kcal mol*^*-*1^. For set S1131, we obtained highest *R*_*p*_ of 0.85 and RMSE of 1.32 *kcal mol*^*-*1^. Finally, for set S8338, our method attained the highest *R*_*p*_ of 0.88 and the lowest RMSE of 0.98 *kcal mol*^*-*1^. The former four methods are representative of advanced geometric-based machine learning methods, while the latter three are mainstream energy-based methods. Comparison results are shown in Table 1 and Figure1. In summary, based on the 10-fold cross-validation, MechPPI achieved overall better performance compared with other methods based on high *R*_*p*_ and lower RMSE.

**Table 1:** Comparison of the *R*_*P*_ of the baseline methods on PPI ΔΔ*G* datasets based on the 10-fold cross-validation results reported in their original papers.

| Method | s4169 |  | s8338 |  | s1131 |  |
| --- | --- | --- | --- | --- | --- | --- |
| | $R_p$ | RMSE | $R_p$ | RMSE | $R_p$ | RMSE |
| **MechPPI** | **0.783±0.03** | **1.11±0.07** | **0.88±0.02** | **0.98±0.06** | **0.85±0.04** | **1.32±0.19** |
| TopNetTree [17] | 0.79 | 1.13 | 0.85 | 1.11 | 0.85 | 1.55 |
| GeoPPI [12] | 0.78 | - | 0.85 | - | 0.85 | - |
| DGCddG [18] | - | - | - | - | 0.84 | 1.30 |
| mCSM-PPI2 [13] | 0.76 | 1.19 | 0.82 | 1.18 | - | - |
| MutaBind2 [14] | 0.74 | - | 0.81 | - | - | - |
| FoldX [9] | 0.27 | - | 0.44 | - | 0.45 | - |
| BeAtMuSic [19] | - | - | - | - | 0.27 | - |

**Figure 1:**
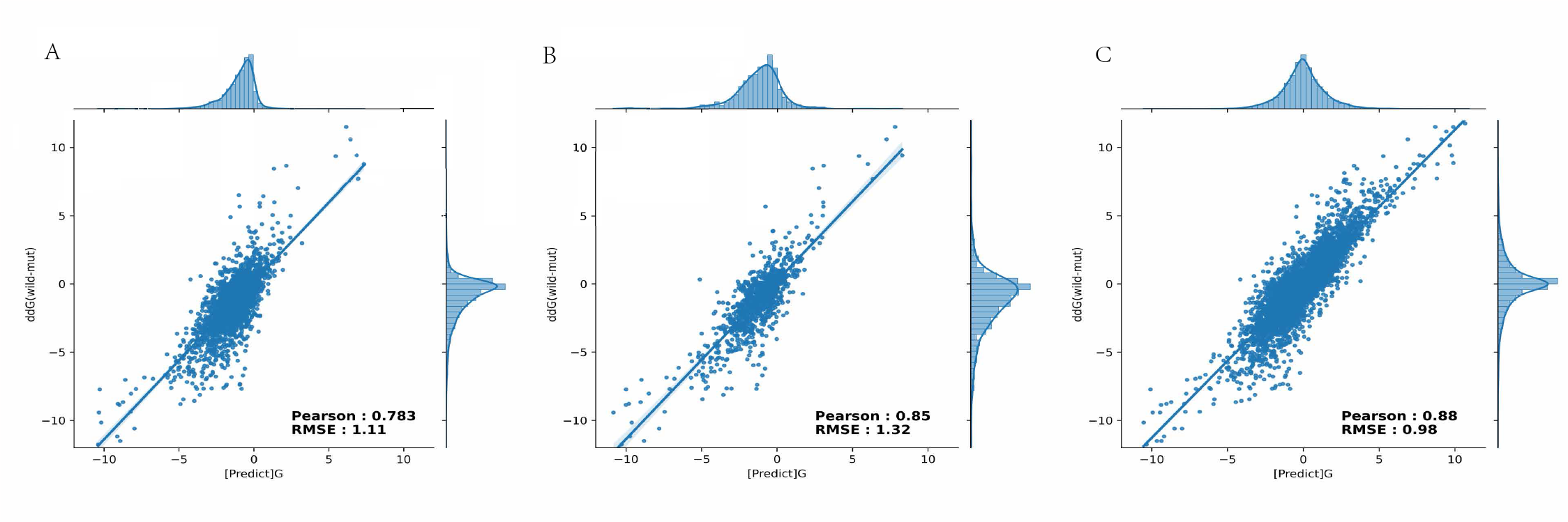
Correlation between predicted and experimental ΔΔ*G* values on s4169 (A), s1131 (B) and s8338 (C)

#### 3.1.2 Protein-level cross-validation

We also performed protein-level cross-validation on several single-point mutation datasets. 10-fold cross-validation is the process of testing the fit of a model to the training set rather than an unseen testing set; thus, a good cross-validation performance does not guarantee the avoidance of overfitting. Protein-level cross-validation can ensure that the data in the testing fold are not leaked into the training folds, this process simply prevents similar protein structures from appearing in the folds of the training set. Based on the Protein-level cross-validation cross-validation, For set S4169, we obtained an average *R*_*p*_ of 0.50 and RMSE of 1.58 *kcal mol*^*-*1^. For set S1131, we obtained *R*_*p*_ of 0.33 and RMSE of 2.23 *kcal mol*^*-*1^. Finally, for set S8338, our method attained the best *R*_*p*_ of 0.71 and RMSE of 1.49 *kcal mol*^*-*1^.Comparison results are shown in Table 2

**Table 2:** Comparison of the *R*_*P*_ of the baseline methods on PPI ΔΔ*G* datasets based on the protein-level cross-validation results reported in their original papers.

| Method | s4169 |  | s8338 |  | s1131 |  |
| --- | --- | --- | --- | --- | --- | --- |
| | $R_p$ | RMSE | $R_p$ | RMSE | $R_p$ | RMSE |
| MechPPI | **0.50**±0.12 | **1.58**±0.42 | **0.71**±0.08 | **1.49**±0.33 | **0.34**±0.19 | **2.23**±0.91 |
| TopNetTree | 0.41 | 1.60 | 0.59 | 1.65 | 0.34 | 2.31 |
| FoldX | 0.27 | 2.73 | 0.44 | 2.73 | 0.46 | 2.18 |
| DGCddG | 0.38 | 1.63 | 0.55 | 1.68 | 0.48 | 2.14 |

**Table 3:** The effectiveness of four mechanism feature groups on subsets with different mechanisms.

| Used Feature Group | Electrostatic Subset |  | Steric subset |  | Hydrophobic subset |  | Hbond subset |  |
| --- | --- | --- | --- | --- | --- | --- | --- | --- |
| Steric feature group | 0.48 ± 0.18 | 1.64 ± 0.42 | **0.59**±0.18 | **1.30**±0.20 | 0.61 ± 0.16 | 1.84 ± 0.44 | 0.62 ± 0.09 | 1.52 ± 0.26 |
| Hydrophobic feature group | 0.37 ± 0.18 | 1.71 ± 0.43 | 0.59 ± 0.18 | 1.28 ± 0.25 | **0.66**±0.16 | **1.76**±0.40 | 0.52 ± 0.11 | 1.72 ± 0.27 |
| Electrostatic feature group | **0.81**±0.16 | **1.02**±0.38 | 0.50 ± 0.25 | 1.41 ± 0.39 | 0.66 ± 0.25 | 1.76 ± 0.25 | 0.62 ± 0.087 | 1.55 ± 0.13 |
| Hbond feature group | 0.66 ± 0.31 | 1.32 ± 0.48 | 0.40 ± 0.24 | 1.50 ± 0.40 | 0.52 ± 0.23 | 2.01 ± 0.47 | 0.57 ± 0.08 | 1.64 ± 0.21 |
| Hydrophobic+Electrostatic+Hbond(No Steric) | 0.81 ± 0.08 | 1.04 ± 0.35 | **0.62**±0.18 | **1.22**±0.37 | 0.71 ± 0.22 | 1.64 ± 0.23 | 0.67 ± 0.09 | 1.44 ± 0.22 |
| Electrostatic+Hbond+Steric(No Hydrophobic) | 0.82 ± 0.10 | 1.02±0.38 | 0.64 ± 0.16 | 1.21 ± 0.24 | **0.70**±0.22 | **1.66**±0.32 | 0.71 ± 0.09 | 1.36 ± 0.23 |
| Hydrophobic+Hbond+Steric(No Electrostatic) | **0.73**±0.14 | **1.19**±0.47 | 0.65 ± 0.16 | 1.21 ± 0.26 | 0.71 ± 0.18 | 1.64 ± 0.32 | 0.71 ± 0.08 | 1.36 ± 0.23 |
| Hydrophobic+Electrostatic+Steric(No Hbond) | 0.78 ± 0.16 | 1.15 ± 0.33 | 0.71±0.13 | 1.10 ± 0.27 | 0.71±0.19 | 1.66 ± 0.25 | **0.69**±0.08 | **1.40**±0.22 |

### 3.2 MechPPI accurately predicts stabilizing, destabilizing and neutral mutation

Even though we can predict the value of ΔΔ*G*, which can precisely reflect the impact of mutation on protein-protein interaction. However, skempi 2.0 include bias and variance in its data, besides, experimental data of skempi 2.0 might be affected by errors. When the value of the free energy change is close to 0 and the associated error is considered, for one single measure the sign of ΔΔ*G* can change from decreasing to increasing and vice versa. Another problem is that the training data are intrinsically non-symmetric and unbalanced, with destabilizing mutations outnumbering stabilizing ones (see Figure 2). This can bias training and testing, affecting the prediction results. The experimental ΔΔ*G* values are affected by uncertainty as measured by standard deviations. Most ΔΔ*G* values are within -0.5 *kcal mol*^*-*1^ to 0.5 *kcal mol*^*-*1^. In order to overcome this problem we describe a new predictor that discriminates between 3 mutation classes: destabilizing mutations (ΔΔ*G <* −0.5 *kcal mol*^*-*1^), stabilizing mutations (ΔΔ*G >*0.5 *kcal mol*^*-*1^) and neutral mutations (−0.5 *<* ΔΔ*G <*0.5 *kcal mol*^*-*1^). We use the Gradient Boosting Tree classifier to predict if the mutation belongs to a stabilizing, destabilizing mutation, or neutral mutation. For s4169, we achieved an average accuracy of 0.80, precision of 0.76, recall of 0.69, and F1 of 0.71. We also get high AUC for stabilizing mutation(0.812), destabilizing mutation(0.802), or neutral mutation(0.780) as Figure 3 showed.

**Figure 2:**
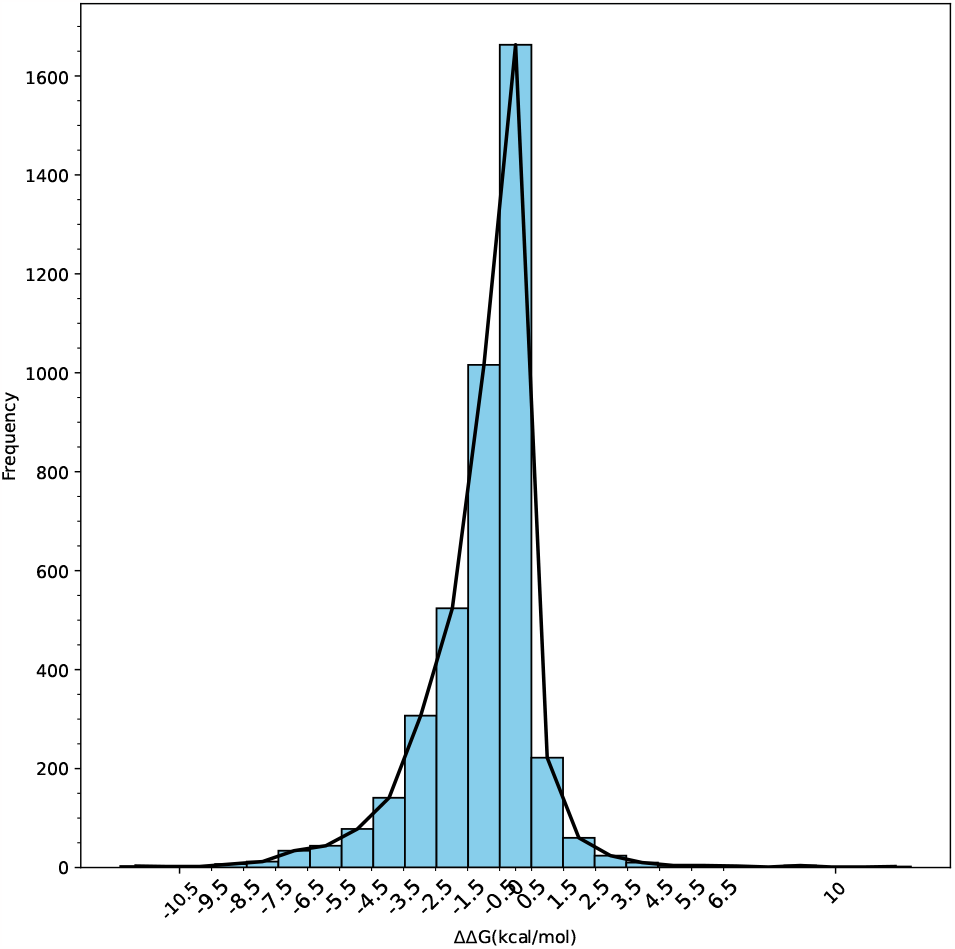
Distribution of ΔΔ*G* in s4169

**Figure 3:**
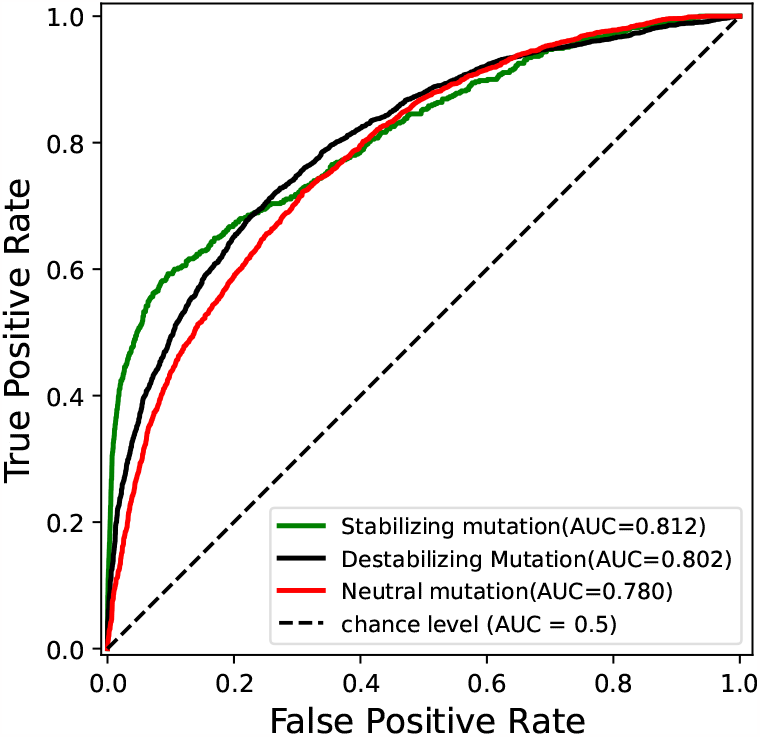
Stabilizing, Destabilizing and Neutral mutation prediction for s4169

### 3.3 The effectiveness of four mechanism feature groups on subsets with different dominant mechanisms

We investigated the efficacy of four distinct mechanism feature groups when applied to different subsets of protein-protein interactions characterized by varying dominant mechanisms in response to mutations. We created a protein-protein interactions dataset that not only includes mutation information but also includes mechanisms underlying mutations by looking through corresponding papers. For example, if a mutation happened, in the paper we found hydrogen bonds disappear due to mutation, and then we would annotate the mutation as a hydrogen bond mechanism. Besides, we also annotated the hydrophobic effect, electrostatic complementarity, and steric effect by looking through papers. In total, we get 952 entries with manually annotated mechanisms, 186 entries include mutations related to hydrophobic effects, 508 entries that mutations related to hydrogen body, 98 entries are related to electrostatic complementarity, and 160 entries are related to steric effects. In order to investigate the efficacy of four distinct mechanism feature groups, we use different combinations of feature groups to predict the binding affinity, each time we remove one type of feature group and use three types of features to predict binding affinity when we drop off specific feature groups and use a subset with similar annotation mechanism, the performance should be the lowest, which can provide information that feature groups are most important for this dataset with specific mechanism. All results are shown in Fiugure3. For the electrostatic subset, when we only use the electrostatic feature group, we get *R*_*p*_ of 0.81, higher than other feature groups. Furthermore, if we remove the electrostatic feature group, we get the lowest *R*_*p*_ of 0.73 compared with the results that removing other feature group. This showed electrostatic feature group can be accurately and effectively used to capture electrostatic complementarity change underlying mutations and produce accurate prediction. The same trend is also found in the steric subset and hydrophobic subset, when only using one feature group, the corresponding feature group performs best, and when removing the corresponding feature group for prediction, the performance is the lowest. However, the hydrogen bond subset did not show a similar trend, it showed our hydrogen bond features might not so sensitive enough for binding affinity change.

### 3.4 MechPPI performance for different mutation types

According to the point mutation distance to the opposite chain, point mutation can be classified as interfacial mutation and noninterfacial mutation. We get a higher average energy change of around 0.86 *kcal mol*^*-*1^ for interfacial mutation and 0.80 *kcal mol*^*-*1^.

According to the locations of the site, mutations could be categorized into five different regions: interior, surface, rim, support and core. In experimental data, mutations at the core or support region have a higher average energy change of around 0.89 *kcal mol*^*-*1^ and 0.83 *kcal mol*^*-*1^, respectively, whereas mutations at the rim or interior region have an average energy change of around 0.74 *kcal mol*^*-*1^ and 0.73, respectively, as shown in Fig. The surface mutations have an average energy change of less than 0.54 *kcal mol*^*-*1^. A possible reason for these patterns is that different mutation regions vary in their accessibility to water; in general, surface, interior, and rim regions have greater access to water than the core and support regions.

We categorized protein complexes as six types: 1) G-proteins, cell cycle, signal transduction, 2) Miscellaneous, 3) Enzyme complexes, 4) Protease-inhibitor, 5) Antibody-antigen, 6) Large protease complexes. Mutations from Protease-inhibitor have a higher average energy change of around 0.93 *kcal mol*^*-*1^ than other catogories. While mutations from Large Protease complexes have a lower average energy change of around 0.66 *kcal mol*^*-*1^. For other protein complexes, energy changes are similar. All the above results are shown in Figure 4

**Figure 4:**
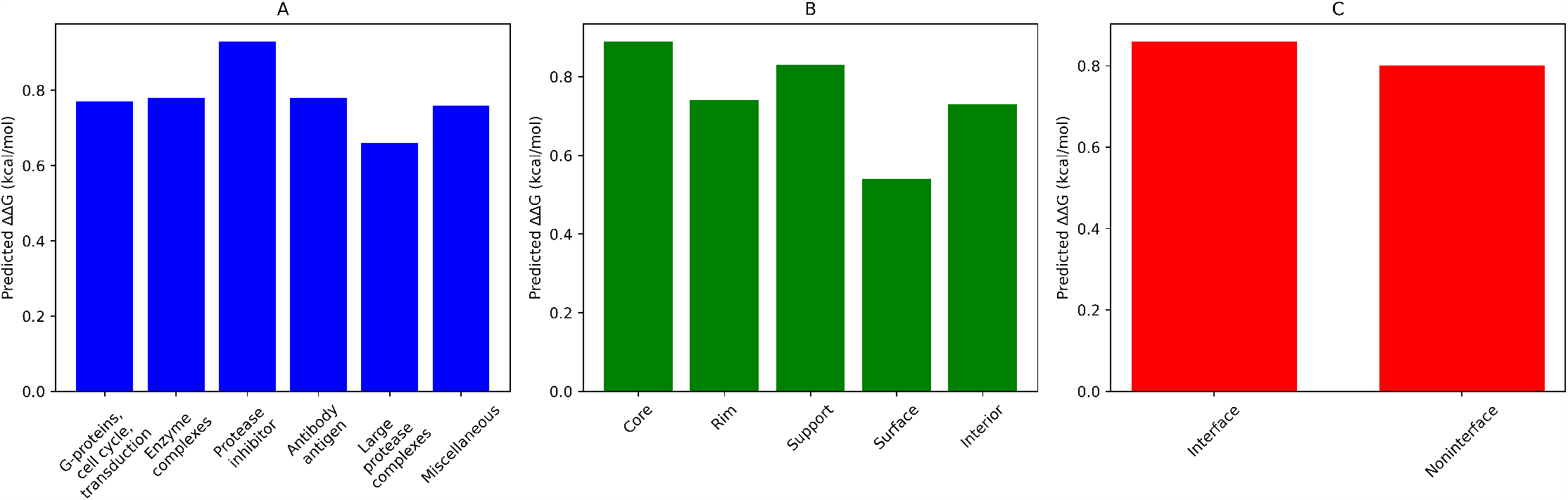
Performance of different mutation types,(A) is prediction performance across different protein categories, (B) is prediction performance across different mutation locations, (C) is prediction performance between interfacial mutation and non-interfacial mutation

## 4 CONCLUSION

MechPPI is a tool that can use potential mechanism features underlying mutation to predict the binding affinity change upon mutation. It can accurately pre dict binding affinity change upon a single mutation, and results demonstrate the potential of MechPPI as a powerful and useful computational tool in protein design and engineering.

